# Ancient genomes reveal long range influence of the site and culture of Tiwanaku

**DOI:** 10.1101/2021.01.22.427554

**Authors:** Danijela Popović, Martyna Molak, Mariusz Ziołkowski, Alexei Vranich, Maciej Sobczyk, Delfor Ulloa Vidaurre, Guido Agresti, Magdalena Skrzypczak, Krzysztof Ginalski, Thiseas Christos Lamnidis, Nathan Nakatsuka, Swapan Mallick, Mateusz Baca

## Abstract

Tiwanaku was a civilization that flourished in the Lake Titicaca Basin (present-day Bolivia) between 500 and 1000 CE. At its apogee, Tiwanaku controlled the lake’s southern shores and influenced certain areas of the Southern Andes. There is a considerable amount of archaeological and anthropological data concerning the Tiwanaku culture; however, our understanding of the population of the site of Tiwanaku is limited. To understand the population dynamics at different stages of the Tiwanaku cultural development, we analyzed 17 low-coverage genomes from individuals dated between 300 and 1500 CE. We found that the population from the Lake Titicaca Basin remained genetically unchanged throughout more than 1200 years, indicating that significant cultural and political changes were not associated with large scale population movements. In contrast, individuals excavated from Tiwanaku’s ritual core were highly heterogeneous, some with genetic ancestry from as far away as the Amazon, supporting the proposition of foreign presence at the site. However, mixed-ancestry individuals’ presence suggests they were local descendants of incomers from afar rather than captives or visiting pilgrims. A number of human offerings from the Akapana Platform dating to ca. 950 CE mark the end of active construction and maintenance of the monumental core and the wane of Tiwanaku culture.

**Significance Statement:** Tiwanaku was an important pre-Inca polity in South America and an example of primary social complexity on par with civilizations in the Indus and Nile river valley. Flourishing between 500 and 1000 CE, Tiwanaku exercised control in the south Titicaca basin and influenced a vast area in southern Peru, Bolivia, and northern Chile. Comprehensive archeological studies provided information about the rise, expansion, and fall of the Tiwanaku culture, but little is known about the monumental site’s population. To address this lacuna, we generated low coverage genomes for 17 individuals, revealing that while the Titicaca basin’s residential population was homogenous, the individuals excavated from the ritual core of Tiwanaku drew their ancestry from distant regions.

## Introduction

The motivations and desires shaping the transition from small-scale societies to settled life – villages, cities – is one of the primary questions in archaeology. Located 3850 meters above sea level near the shores of Lake Titicaca, Tiwanaku represents the unlikeliest case of the spontaneous and primary rise of social complexity on par with the handful of other locations on earth (Fig. 1A). For almost half a millennium (500-1000 CE), Tiwanaku was one of the most influential centers in the Southern Andes (Fig. 1B).

**Figure 1.**
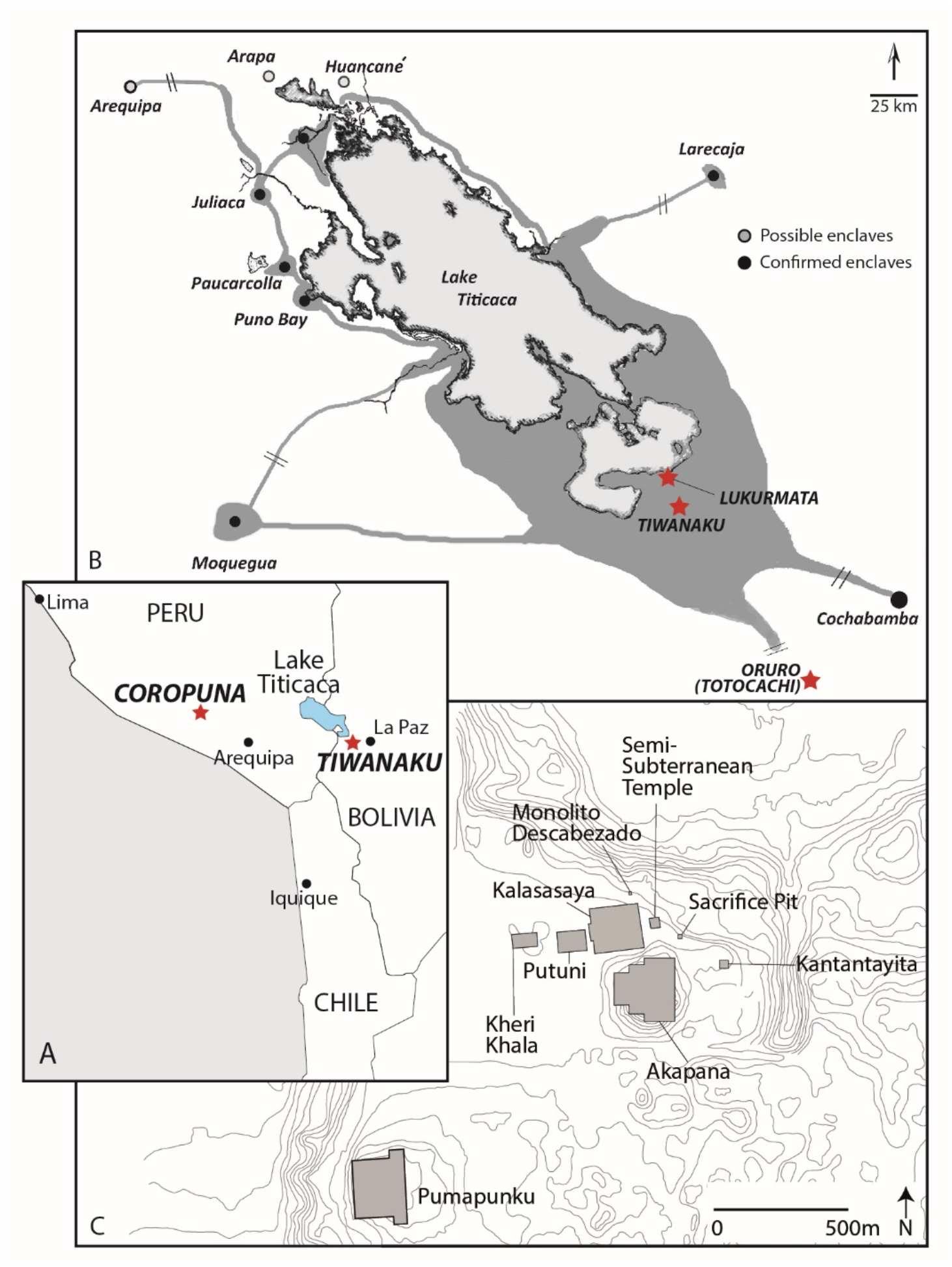
Provenience of the studied individuals. A) Map of a part of South America with Tiwanaku and Coropuna locations from which the individuals were sampled are indicated with red stars, as well as Lake Titicaca, major cities, the modern country borders and the coastline. B) Extension of Tiwanaku influence with main sites and locations from where TIW, LUK and ORU individuals were sampled for this study are indicated with red stars (redrawn from Stanish (2003); map to scale around the lake, not to scale in outlying regions). C) Location of the monuments at the site of Tiwanaku (contours redrawn from Kolata (2003)).

The heavily deteriorated monuments of Tiwanaku are well known and described in detail from the initial years of the Spanish Conquest (Ponce Sanginés, 1995). More than one hundred years of archeological research reveals how cultural and demographic changes in the Lake Titicaca Basin precedes Tiwanaku’s emergence as the primary ritual center around 500 CE. The south-eastern Basin enjoys a milder climate than other parts of the Altiplano. Densely inhabited from at least the Early Middle Formative Period (1500 – 100 BCE), the three southern valleys – Desaguadero, Tiwanaku, and Katari, hosted several pre-Tiwanaku cultures. With the emergence of the Tiwanaku, demographic changes took place, with some settlements reducing in size as people were drawn to this monumental site. For example, Khonko Wankane, a large site in the Desaguadero valley, became mostly abandoned by the end of the Late Formative period (ca. 500 CE); conversely, Lukurmata remained an important settlement in the adjacent Katari Valley (Janusek, 2004a).

Like the other Andean cultures, Tiwanaku ritual practices forged a degree of political unity among diverse groups (Janusek, 2004b). The ritual core of the city of Tiwanaku was designed according to beliefs about unity between cosmological, mythical space, and physical space (Vranich, 2006). The earliest monument at the site, the Semi-Subterranean Temple, was complemented in the 3th - 5th century CE with adjacent Kalasasaya Platform and Kheri Khala complex. Around 1km away is the Pumapunku Temple Complex and several additional minor monuments further to the south (Vranich, 2006) (Fig. 1C). The next important phase in the city transformation started around 600 CE with the construction of the Akapana Platform. A few centuries later, renovations to the existing building and the construction of new structures such as the Putuni and Kantatayita would create the site one can presently see. Between and around these monuments were plaza areas and residential compounds.

Construction of the largest building at the site, the Akapana Platform, began in the middle of the 7th century and was a conglomeration of reused stone (Vranich, 2001). Visible from several kilometers away, this imposing platform consisted of seven stepped terraces. At the platform’s base are numerous humans remains along with camelid bones and ceramic sherds (Manzanilla and Woodard, 1990; Escalante, 2006; 2007; 2008). These remains were interpreted by the excavators as ritual offerings placed in soil accumulated above the base of the Akapana Platform (Blom & Janusek, 2004).

Tiwanaku culture was pre-literate, which means that archaeological research is the only source of primary information. The last century of research provides enough information to address fundamental issues such as layout, size, and distribution of the buildings; nevertheless, a fundamental question remains about the site’s population. Scholarly opinions range from a densely occupied diverse city to a near-empty ceremonial central that cyclically pulsated with life from seasonal pilgrims (Janusek, 2013). Another critical issue is the ancestry of the human offerings found associated with the ritual platforms. A combination of archaeological and genetic evidence provides insight into the types of people buried on and next to Tiwanaku’s ceremonial structures. For this research, we examine genome-wide information for 13 ancient individuals from Lake Titicaca region in Bolivia associated with the Tiwanaku culture (500-1000 CE), and four from the region of Coropuna volcano in southern Peru associated with Wari (500-1000 CE) and Inca cultures (ca. 1400-1540 CE, in this region). We analyze the genetic make-up of these groups, compare their affinities to other ancient and modern populations, and determine the genetic ancestry of the human offerings at the site.

### Ethics statement

All samples for the individuals included in this study were obtained under permissions from the local authorities.

Individuals from Bolivia were sampled for analyses under permission from La Unidad de Arqueologia y Museos (UDAM) Ministerio de Culturas y Turismo Bolivia no. and 052/2016 and 086/2016. Samples from Peruvian individuals were collected under permissions granted by Peruvian Ministerio de Cultura (formerly Instituto Nacional de Cultura). Results obtained in this study will be provided to authorities in Centro de Investigaciones Arqueológicas, Antropológicas y Administración de Tiwanaku (CIAAAT). We will collaborate with CIAAAT to share our research findings with the local community.

## Results and Discussion

We screened 93 specimens sampled from pre-Columbian sites near Lake Titicaca in Bolivia and Southern Peru (Dataset S1*A*; S1*B*) for DNA preservation. To increase the very low content of endogenous DNA we used various approaches, including whole genome capture (Carpenter et al., 2013), pre-digestion (Damgaard et al., 2015), and, in the case of teeth, drilling only the outer layer of the roots (cementum) (Higgins et al., 2013). Consequently, we generated low-coverage genome sequences for 18 individuals with a depth of coverage ranging from 0.15x to 2.56x. The sequencing data showed deamination pattern at 5’ and 3’ ends and mean fragment length characteristic for ancient DNA (Dataset S1*A*). Authenticity of the data was corroborated by lack of detectable contamination in the nuclear DNA and low estimates for the mitochondrial DNA contamination and lack of the heterozygosity on the X chromosome in male specimens (below 5% for all estimations) (Dataset S1*A*). One individual (TW098) did not pass the quality control thresholds for ancient DNA authenticity, with the nuclear contamination estimated at around 9%, and was removed from further analyses. We found slightly higher contamination estimated for X chromosome (5,5%) in case of TW059 individual but considering that it was based on limited number of SNPs and all other contamination estimates were below the threshold we decided to keep this individual in downstream analyses. Dating of the ancient individuals based on archaeological context and stratigraphy was confirmed by direct radiocarbon dating, thus strengthening their chronological setting (Table 1; Dataset S1*A*; Fig. S1).

**Table 1.**
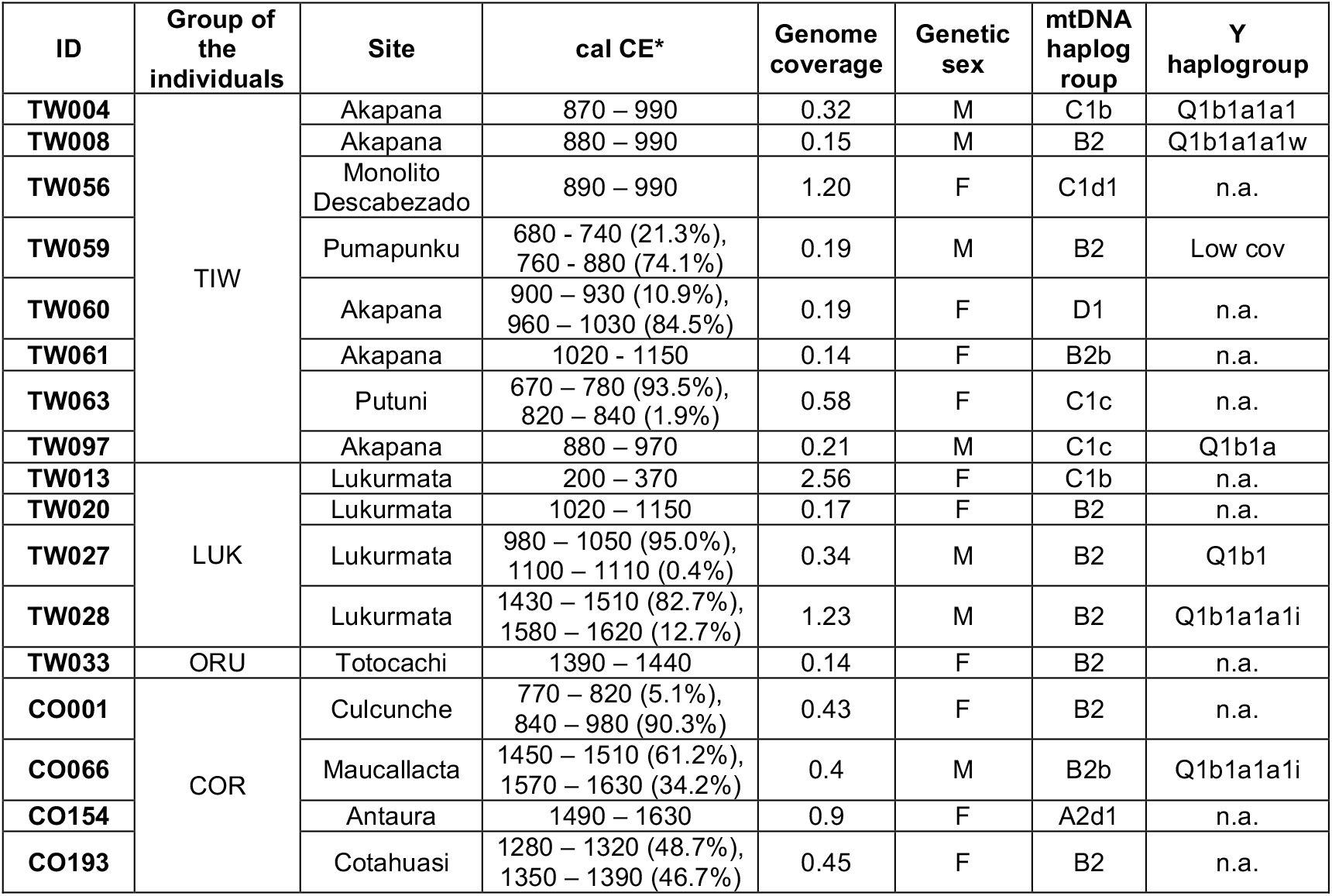
Radiocarbon dates and main genetic indices of analyzed individuals.

Our dataset contains 13 individuals from the Lake Titicaca basin covering the period between ca. 300 and 1500 CE. The residential group from outside the Tiwanaku ritual core site consisted of five individuals. Four of them originated from the site of Lukurmata in nearby Katari Valley (TW013, TW020, TW027, TW028; thereafter LUK). Individual TW033 (thereafter ORU) was excavated in Totocachi, Oruro, separated from Katari Valley by two hundred kilometers.

Radiocarbon dating revealed that within the LUK group, there were individuals representing three periods. Individual TW013 was dated to ca. 300 CE and predates the Katari Valley incorporation in the later Tiwanaku polity. Two other individuals (TW020 and TW027) date to between ca. 980 and 1100 CE, the period witnessing the decline and abandonment of the Tiwanaku site. Finally, TW028 (1470 CE) falls into the period of the occupation of the area by the Inca Empire. The ORU individual is dated after the abandonment of Tiwanaku site and perhaps already represents the later cultures. The other eight individuals originated from five different locations within the ritual core of the Tiwanaku site (TIW): the base and revetments of the Akapana Platform (TW060, TW061), the Pumapunku Platform (TW059), south side of the Putuni Platform (TW063), the offering pit between the Semi-Subterranean Temple and the Akapana Platform (TW097), and an area of midden adjacent to a monolith (Monolito Descabezado) along the north-eastern corner of the Kalasasaya Platform (TW056) (Fig. 1C). Two individuals do not have a precise provenience (TW004, TW008), but they are suspected of having come from excavations at the Akapana Platform. Two individuals (TW059 and TW063) lived in a period of strong Tiwanaku influence in the basin and southern Andes (700 - 800 CE), while others date approximately to 950 CE, a period considered to be one of decline.

In addition to the individuals directly associated with the Tiwanaku culture, genomic data was generated for four individuals from the Late Intermediate Period (LIP; 1000–1450 CE) to Late Horizon (LH; 1400–1540 CE in this region) sites surrounding Coropuna volcano in southern Peru (thereafter COR) (Fig. 1A; Table 1; Dataset S1A). This region was an important ritual and pilgrimage center, during, but also prior to, the Inca period, with traces of Wari and Tiwanaku influence (Tunia, 2005; Ziolkowski & Tunia, 2005). Individual CO001, dating to 920 CE, comes from a pastoralist burial at Culcunche and most probably represents a local population that later hosted the ceremonial center of Maucallacta. Individual CO066 (1500 CE) originated from the Maucallacta while individual CO154 (1560 CE) is from Antaura, settlement located nearby Maucallacta. The last individual (CO193) was a mummy from the nearby Cotahuasi Valley, dating at 1320 CE.

We first performed qualitative assessment of the genetic affinities of the studied individuals using Principal Component Analysis (PCA) and unsupervised ADMIXTURE analyses. Principal Components (PCs) were computed using present-day individuals without European ancestry from South America (SI Text). Genomic datasets of the present-day individuals were generated using different techniques and their final intersection resulted with 199,175 common SNPs (Dataset S1*C*). Ancient individuals from this study and other available ancient South American individuals were projected onto the computed PCs (Fig. 2A; 2B). PCA showed that the majority of individuals cluster within their populations. A split between the Amazonian and the Andean populations was also demonstrated, corroborating previously published results (Barbieri et al., 2019; Gnecchi-Ruscone et al., 2019). COR individuals from this study clustered together with other ancient Peruvian individuals, closest to the Southern Peruvian Highlands (Laramate) and Southern Peruvian Coast (Ullujaya and Palpa) (Nakatsuka et al., 2020a).

**Figure 2.**
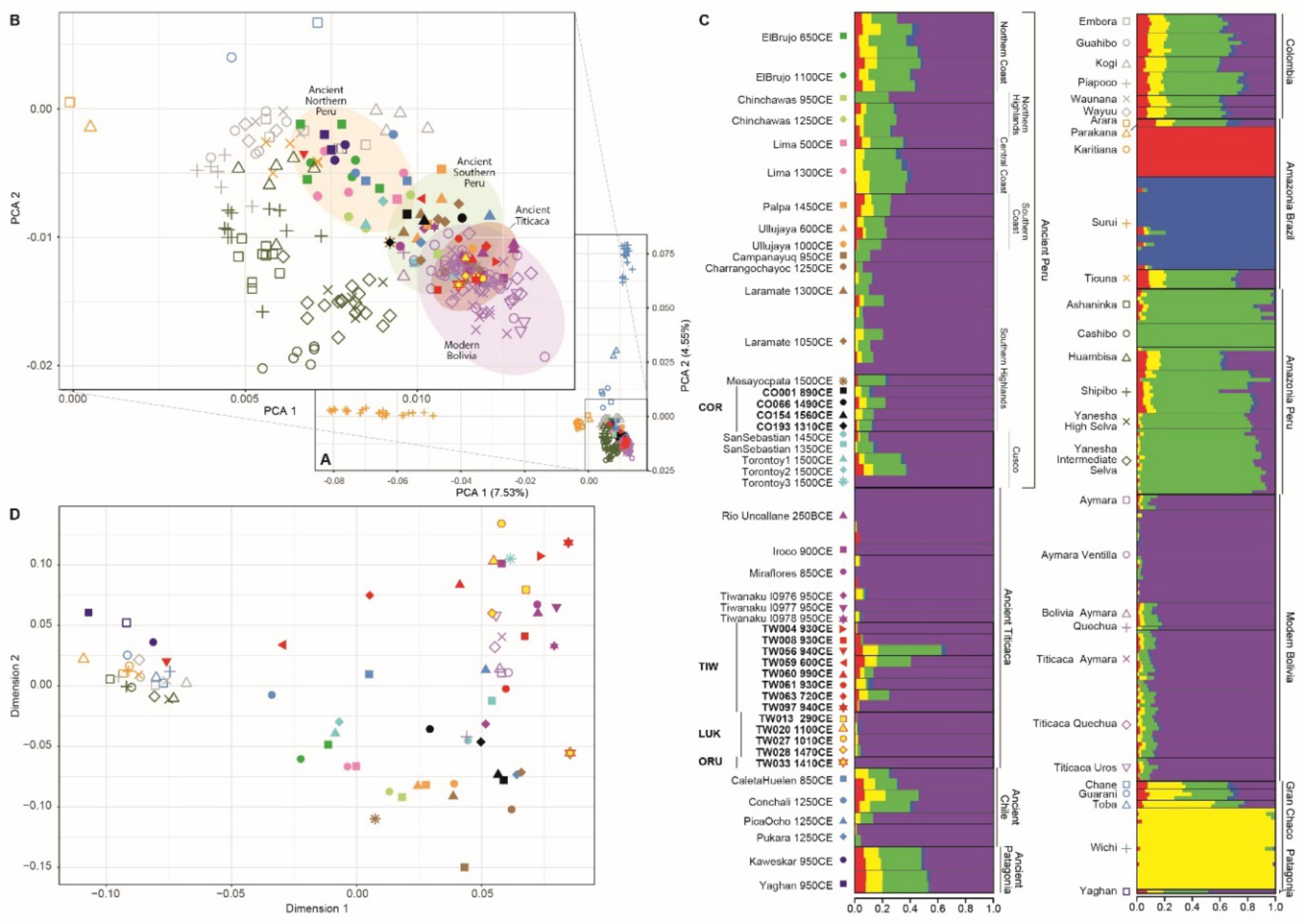
Comparison of the SNP-based genomic variation of the analyzed individuals and previously published ancient and present-day South American genomes. (A) PCA plot of 217 present-day South American individuals (no-fill labels) with ancient individuals projected (filled labels) (B) Zoom into the most relevant section of the plot (C) Admixture plot for K=5. (D) Multi-Dimensional Scaling (MDS) plot for the *f*_*3*_ statistics (1-*f*_*3*_*(IND, test, Mbuti)*) matrix.

Present-day Bolivians made up a tight cluster with ancient individuals from Lake Titicaca Basin. Two of the TIW individuals, however, fell notably outside this ancient Titicaca cluster. TW056 was projected within the dispersed cluster of modern Amazonian, Columbian and ancient individuals from the Northern Peruvian Coast, whereas TW059 was projected between the ancient Northern and Southern Peruvian group (Fig. 2A; 2B). Unsupervised ADMIXTURE analysis showed that considering K=5 (the lowest CV error, Dataset S1*D*; Fig. 2C) two of the ancestral components were dominant and characteristic for two Amazonian populations from Brazil - Karitiana and Surui (red and blue, respectively). One component (yellow) was characteristic for Wichi, population from the Gran Chaco (Northern Argentina). The last two components were dominant in the Peruvian Amazonian (green) and in the Andean populations (violet).

LUK individuals, together with other ancient Lake Titicaca individuals (ORU, Rio Uncallane, Iroco and Miraflores) trace their ancestry mostly to a single (violet) component (>95%). Notably, a present-day high-altitude Bolivian population from Ventilla (Lindo et al., 2018) shares this almost exclusively Andean (violet) genetic make-up, distinguishing them from other present-day Bolivians. On the other hand, the majority of the TIW individuals show a higher proportion of the Amazonian (green) component, ranging from 2% to 46% and reaching the maximum in TW056, TW059 and TW063. The exceptions are TW004 and TW0097 individuals comprising the violet component almost exclusively (Fig. 2C).

To investigate the genetic makeup of the studied individuals more deeply we computed outgroup *f*_*3-*_ and *f*_*4-*_ statistics. In all calculations Mbuti, a hunter-gatherer population from the Central Africa was used as an ‘outgroup’. We performed exhaustive calculations of *f*_*4*_-statistics in the form *f*_*4*_*(Mbuti, Population; Ind1, Ind2)* and *f*_*4*_*(Mbuti, Ind; Group1, Group2)* where *Ind* were individuals from this study while *Population* or *Group* were a population or the group of ancient individuals from the same region and period as defined in Nakatsuka et al., (2020a) or present-day populations. Significantly negative values of Z-score (Z > |3|) suggest that the studied *Ind1* shared more alleles with test *Population* than with *Ind2* (Dataset S2*A*) or with *Group1* than with *Group2* (Dataset S2*B*). Shared genetic drift was measured between each of the 17 studied individuals from this study (*‘Ind’*) and all other individuals/populations (*‘Test’*) from the dataset in the form *f*_*3*_*(Ind, Test, Mbuti)*. We generated a multi-dimensional scaling (MDS) plot and a neighbor-joining tree based on outgroup *f*_*3*_-statistics (Fig. 2D; Fig. 3). All LUK individuals showed the highest affinities with each other or with other ancient individuals from the Lake Titicaca area (Fig. 2D; Fig. 3).

**Figure 3.**
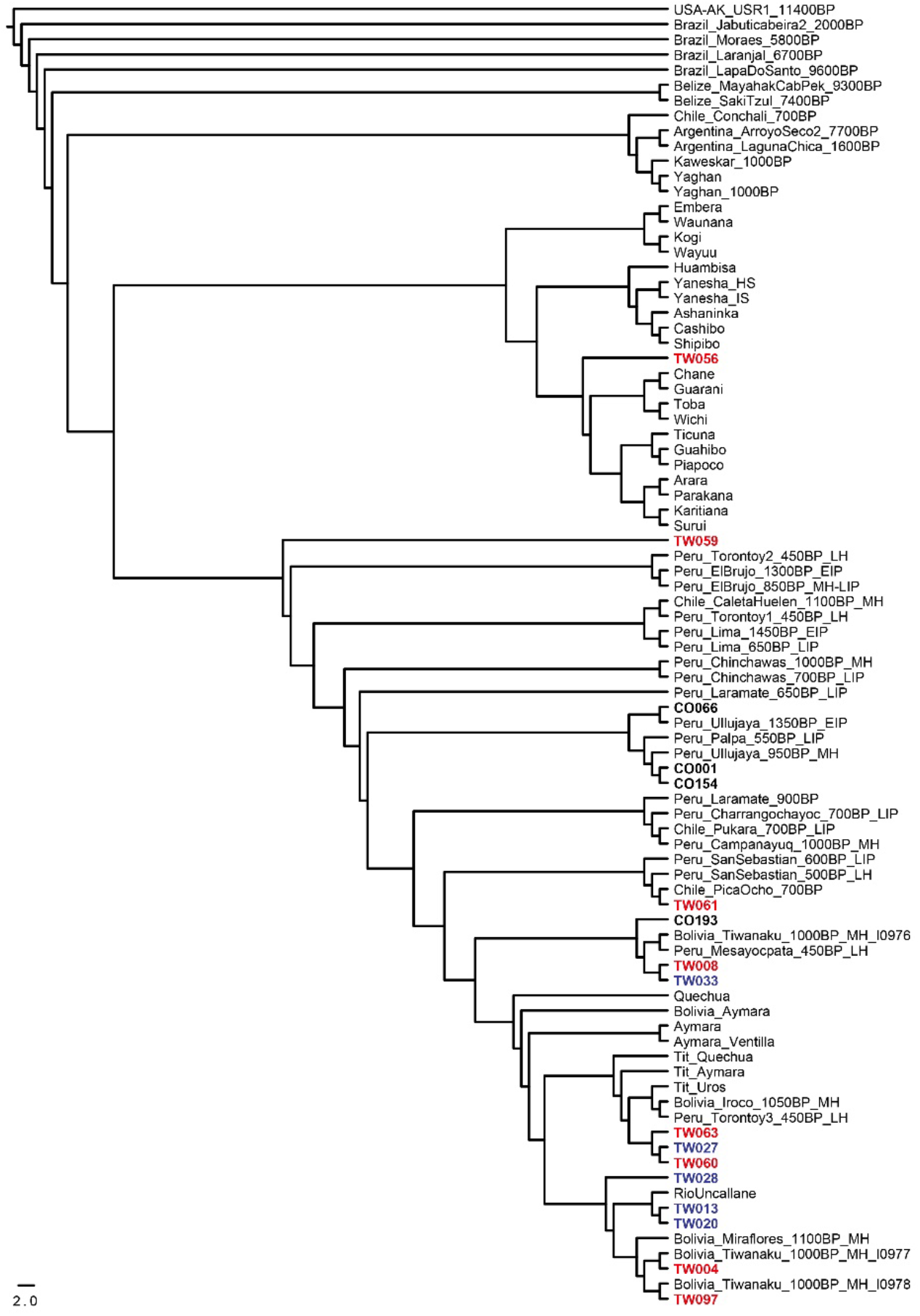
Neighbor Joining Tree based on inverted outgroup-*f3* statistics (1/*f*_*3*_*(IND, test, Mbuti))* visualized using FigTree v1.4.2. Individuals from the current study in bold. TIW in red, LUK and ORU in blue.

Homogeneity of the Lukurmata individuals was tested using *f*_*4*_-statistics in the form *f*_*4*_*(Mbuti, Test; LUKind1, LUKind2)* where *LUKind1* and *LUKind2* were possible pairs of Lukurmata individuals iterating over groups formed of ancient individuals from the same period and region as defined in Nakatsuka et al. (2020a) (Dataset S2*C*). None of these statistics, nor either in the form *f*_*4*_*(Mbuti, Test; LUK, ancientTiticaca)*, were significant showing that none of the Lukurmata individuals shares significantly more alleles with either *Test* group than with either individual from the Lukurmata or with the group of the ancient individuals from the Titicaca area *(RioUncallane, Bolivia_Miraflores_1100BP_MH, Bolivia_Iroco_1050BP_MH)* (Dataset S2*D*). To assess this with higher resolution, we used qpWave which showed that ancestry of all Lukurmata individual is well explained by a single wave from ancient Titicaca (Dataset S2*A*). This shows that Lukurmata population is genetically homogeneous to the limits of our resolution.

Using the same strategy, we found that the group of individuals from the Tiwanaku ritual core is very heterogeneous and various patterns of ancestry could be recognized (Dataset S2; S3). Individuals TW004, TW008, TW060, TW097 as well as published Akapana individuals *Bolivia_Tiwanaku_1000BP_I0977* and *Bolivia_Tiwanaku_1000BP_I0978* (Nakatsuka et al., 2020a) could be modeled with ancient Titicaca as the only source of ancestry. However, individuals TW059, TW061, and TW063 had more complex ancestries. Individual TW059 shows strong evidence of admixture, as this individual cannot be modeled as a single source of ancestry and instead requires at least two sources. The additional source of ancestry is likely Amazonian as only models with ancient Titicaca and Amazonian or Gran Chaco sources fit the data (Dataset S3*C*). Individual TW061’s very low genomic coverage limits the analytical resolution and hence many models cannot be rejected. One of the fitting *f*_*4*_-statistics models indicates solely ancient Titicaca ancestry, but the outgroup *f*_*3*_-statistic demonstrated large amount of shared drift with *Chile_Pukara_700BP_LIP* and *Chile_PicaOcho_700BP* (Fig. 3; Fig. S3), suggesting this individual might have North Chile-related ancestry. Individual TW063 cannot be modeled as a deriving its ancestry from a single source, but the additional source is unclear, because models including ancient Titicaca with several other sources all fit the data (p>0.05) (Datasets S2B; S3C). Individual *Bolivia_Tiwanaku_1000BP_I0976* could be modeled as a mixture of ancient Titicaca and Southern Peru Highlands (Dataset S3*C*), consistent with Nakatsuka et al. (2020a), but we found also a signal of North Chile ancestry (Dataset S3*B*). However, outgroup *f*_*3*_-statistics showed closeness with Peruvian groups (Fig. 2D; Fig. 3). The most outstanding individual was TW056 with almost exclusively Amazonian-related ancestry indicated by both *f*_*4*_-statistics and qpWave (Datasets S3B). Outgroup *f*_*3*_-statistics indicated the Peruvian Amazonian population of Cashibo as sharing the most genetic drift with this individual (Fig. S3).

Overall, our results indicate that the population of the Titicaca basin was rather homogenous beginning at least at the Formative Period (1800–900 BCE) up until the arrival of the Europeans. This is well illustrated for the Lukurmata site for which we have the widest time transect consisting of four individuals (TW013, TW020, TW027, TW028) from between ca. 300 and 1500 CE, who exhibit genetic similarity and indicate that over at least twelve centuries no major genetic turnovers took place despite significant cultural and political changes. This supports the hypothesis proposed by Bermann (2003) according to which formation of the Tiwanaku state did not influence residential areas.

Earlier genetic studies of present-day South American populations, both using uniparental markers (Fuselli et al., 2003) and genomic data (Gnecchi-Ruscone et al., 2019; Harris et al., 2018), suggested that both pre-contact and present-day populations from the Central Andes were homogenous and characterized by relatively high levels of gene flow. In contrast we find little overlap between pre-contact individuals from Central and Southern Peru, and those from the Lake Titicaca Basin (Fig. 2; Fig. 3). The explanation for that may be the fact that these two regions remained under the influence of distinct polities, Wari and Tiwanaku, respectively. Our results corroborate the findings of Nakatsuka et al., (2020a) who also suggested homogeneity within distinct regions of the Andes during the last ca. 2000 years. However, the contacts between various parts of the Andes were not completely absent, as shown by the single individual from the Totocachi site (TW033) with genetic affinity to individuals from the South Peruvian Andes (Dataset S4*D*). Within the context of genetic homogeneity of the Bolivian Altiplano populations, individuals from the Tiwanaku ritual core stand out with their striking heterogeneity. We found that several individuals reveal close affinities to the Titicaca basin population, while others showed distinct ancestry. Previous studies based on strontium isotopes (Knudson et al., 2004) and cranial modifications (Blom, 2005) proposed the non-local origin of some of the excavated individuals. Artifacts such as jaguar canines (Janusek, 2004a), tropical botanical remains (Manzanilla, 1992), and clear evidence of the consumption of lowland hallucinogenic substances (Berenguer, 1987; Browman, 1981) indicate contacts with the Amazon. Our finding of the individuals tracing their ancestry to remote areas like Amazonia or Gran Chaco confirms that the site attracted people from outside the Titicaca basin and that contact between these cultures across the Andes was not limited to the cultural exchange.

It is important to note that the archaeological context of the Tiwanaku human remains for this study is not residential, as are the individuals from Lukurmata. Instead, they are considered ritual offerings associated with the ceremonial buildings. Previous analysis of individuals along the base of the Akapana found evidence of forceful dismemberment at the time of or soon after death (Blom & Janusek, 2004). A shallow deposit between the Semi-Subterranean Temple and the Akapana contained a group of 16 individuals, several of which showing signs of injuries associated with fighting, along with blunt force trauma to the back of the head before being deposited (Verano, 2013). There is debate about the identities and provenience of these individuals and whether they could be captives from long-distance raiding and organized warfare.

Our results lean towards a more complex hypothesis. The genetic testing of the individual to the north of the Kalasasaya (TW056), whose position and context also supports sacrifice, indicates Amazonian origin. TW097, a loose tooth found near the Akapana in association with an individual placed face down with arms bound behind the back (Vranich and Koons 2006), in turn, presented clear local ancestry. The fragmented individuals from the Akapana (TW004, TW008 and TW060) also show local ancestry, whereas a complete individual from the Pumapunku (TW059) and a solitary skull from a disturbed context at the Putuni (TW063) presented a mixture of local and remote ancestry, which may suggest that they were descendants of migrants who settled at Tiwanaku rather than captives brought by military raids from afar. Overall, our results indicate that the practice of human sacrifice in the Tiwanaku was complex and included locals and foreigners who were brought as captives or came voluntarily.

One remarkable finding is the dating of human offerings. Most of human remains from the Akapana investigated here, and earlier (Llamas et al., 2016), date to the middle of the 10th century CE (Table 1; SI Text). There are other documented cases where such an intensification of human sacrifice indicates a society in crisis, grasping for a solution to an environmental catastrophe (Prieto 2019). The manner and process of the abandonment of the site of Tiwanaku is a subject of debate, but some scholars consider the last portion of the 10th century as the start of a gradual or sharp slide towards the site becoming abandoned (Owens, 2005). Others ascribe the end of Tiwanaku to a long-term drought which started ca. 1100 CE, collapsing the raised field agricultural system (Kolata, 1993; 2003); others find this explanation too environmentally deterministic and point to little evidence of Tiwanaku presence post 1000 CE, even while the raised fields continue for several additional centuries (Erickson, 1999). Our data marks the 10th century as the beginning of the end of the site of Tiwanaku. The intensified offering events last about a century, after which offerings and other evidence of organized activity cease. Undamaged, the primary iconography of Tiwanaku remain standing until the post-contact period. This combination of evidence suggests a structural change at the start of the 10th century, followed by a century of diminishing interest in the site.

## Materials and Methods

### DNA extraction and library preparation

Samples from a total of 93 individuals from modern day Bolivia and Peru were selected for genetic analyses. The human remains originated from archeological sites with contexts and chronologies of Tiwanaku or Inca cultures (Dataset S1B). All work with the anthropological material, DNA extraction and library preparation were performed in a dedicated ancient DNA facility at the University of Warsaw’s Centre of New Technologies and all precautions to avoid any contamination with modern DNA were taken. DNA was extracted from teeth or bone, converted into double-indexed sequencing libraries and sequenced on Illumina platforms (NextSeq500, HiSeq4000 and/or NovaSeq6000) (SI Text).

### Data processing

Adapter sequences were trimmed and paired-end reads were collapsed using AdapterRemoval2 (Schubert et al., 2016). Sequencing reads were mapped to human reference genome h37db5 applying default parameters of the BWA mem algorithm (Li & Durbin, 2010). Samtools (Li et al., 2009) were used to remove duplications and only reads longer than 30bp and with mapping quality over 30 were used in subsequent analyses. To minimize impact of deamination on the potential genotyping errors we trimmed 7bp from both ends of all reads using the trimBam tool from bamUtils (Jun et al., 2015).

### Authentication

We used mapDamage 2.0 (Jónsson et al., 2013) to assess damage and fragmentation patterns of the obtained sequencing reads. Contamination from present-day human mtDNA was checked using schmutzi (Renaud et al., 2015) and contamMix (Fu et al., 2013). Nuclear DNA contamination was assessed using ContamLD (Nakatsuka et al., 2020b). Additionally, for male individuals, we investigated nuclear contamination based on polymorphic sites on the X-chromosome using ANGSD (Korneliussen et al., 2014).

### Sex determination

Genetic sex of the analyzed individuals was determined by calculating the ratio of sequence reads aligning to X and Y chromosomes as it was described by Skoglund et al. (2013) as well as comparing the coverage on the X- and Y-chromosome with the coverage on autosomal chromosomes as it was described by Lamnidis et al. (2018).

### Genotyping and datasets

The genotypes were called choosing one random read for every SNP from the 1240K SNP list (Haak et al., 2015) using the script pileupCaller–a part of sequenceTools (https://github.com/stschiff/sequenceTools). The data were merged with the available modern and ancient genome data from South America (de la Fuente et al., 2018; Gnecchi-Ruscone et al., 2019; Lindo et al., 2018; Nakatsuka et al., 2020a; Posth et al., 2018; Reich et al., 2012). In downstream analyses we used published ancient genomes which possessed more than 20,000 SNPs intersected and were dated to within the last 2000 years.

### Genetic affinities

Principal Component Analysis (PCA) was computed using smartpca script from EIGENSOFT package (Patterson et al., 2006) and *lsqproject=YES* and *shrinkmode=YES* options. Ancient individuals were projected on the PC plot computed with modern un-admixed South American individuals (SI Text; Dataset S1*C*). To investigate the genetic structure of the Tiwanaku populations an unsupervised admixture analysis was performed using software ADMIXTURE (Alexander et al., 2009). Prior to the admixture analysis genotypes were pruned for minor allele frequency below 0.01 and linkage disequilibrium with a window size of 200, a step size of 5 and an R2 threshold of 0.5 using plink (Purcell et al., 2007). Five replicates were done for each K (K=2 to K=15) and the optimal K was chosen based on the lowest cross-validation error (Dataset S1*D*). Outgroup *f*_*3*_-statistics were estimated with qp3pop from ADMIXTOOLS (Patterson et al., 2012). We created matrix of the output *f*_*3*_-statistics values between all pairs of individuals/populations from the dataset. The obtained values were transformed into distances by subtracting the values from 1 and generated MDS plot using cmdscale option in R. Neighbor joining tree was generated using PHYLIP software (Felsenstein, 1993) and individual USA_USR1_AncientBeringian_1140BP.SG (Moreno-Mayar et al., 2014) as an outgroup. To compute *f*_*4*_-statistics, qpWave and qpAdm we used admixR package (Petr et al., 2019) applying option *‘transversion only’*.

## Data availability

BAM files from this study are available from the European Nucleotide Archive under accession number: PRJEB41550.

## Acknowledgments

We are grateful to Centro de Investigaciones Arqueológicas, Antropológicas y Administración de Tiwanaku (CIAAAT) for access to human remains stored in warehouse. We thank Elizabeth Arratia in charge of the warehouses of the CIAAAT and Analy Quiroga for help in the location of the individuals stored in CIAAAT. We are grateful Vanessa Jiminez for providing information about excavation at Monolito Descabezado and Dominika Sieczkowska for help with mummies in Arequipa. This work was financially supported by grants 2014/15/D/NZ8/00285 and 2017/01/X/NZ8/00410 from the National Science Centre, Poland. M.S. was supported by grant 2015/17/D/NZ2/03711 from the National Science Centre, Poland. N.N. is supported by an NIGMS (GM007753) fellowship. NGS was performed thanks to Next Generation Sequencing Core Facility CeNT UW, using NextSeq 500 and NovaSeq 6000 platforms financed by Polish Ministry of Science and Higher Education (decision no. 6817/IA/SP/2018 of 2018-04-10). The Coropuna individuals were collected within the framework of the fieldworks of the Archaeological Project “Condesuyos” financed with the grant 2011/01/m/HS3/03432 from the National Science Centre (NCN), Poland.

