## Supplementary Information Text for "Ancient genomes reveal long range influence of the site and culture of Tiwanaku"

[1] Centre of New Technologies, University of Warsaw, S. Banacha 2c, 02-097 Warsaw, Poland
[2] Museum and Institute of Zoology, Polish Academy of Sciences, Wilcza 64, 00-679 Warsaw, Poland
[3] Centre for Andean Studies, University of Warsaw, Krakowskie Przedmieście 26/28, 00-927 Warsaw, Poland
[4] Department of Anthropology, University of Texas, San Antonio. College of Liberal and Fine Arts. One UTSA Circle San Antonio, TX 78249-1644
[5] Unit of Archeology and Museums - Vice-Ministry of Interculturality. Tiahuanaco Street No93 at the corner of Federico Suazo. Box 4856. La Paz, Bolivia
[6] Department of Archaeogenetics, Max Planck Institute for the Science of Human History, Kahlaische Straße 10, 07745 Jena, Germany
[7] Department of Genetics, Harvard Medical School, Boston, Massachusetts 02115, USA
[8] Harvard-MIT Division of Health Sciences and Technology, Boston, MA 02115, USA
[9] Howard Hughes Medical Institute, Harvard Medical School, Boston, MA 02446, USA
[10]Broad Institute of Harvard and MIT, Cambridge, MA 02142, USA

*corresponding authors:
* Danijela Popović; Mateusz Baca

**This PDF file includes:**

Supplementary Information Text
Figures S1 to S3

**Other supplementary materials for this manuscript include the following:**

Datasets S1 to S3

### Supplementary Information Text

### 1.ARCHAEOLOGICAL SITE INFORMATION

#### Bolivia

Samples from individuals from Bolivia were obtained under permission from La Unidad de Arqueologia y Museos (UDAM) Ministerio de Culturas y Turismo no. and 052/2016 and 086/2016.

#### Tiwanaku

The highland sprawling monumental site of Tiwanaku (16°33'22"S, 68°40'11"W) is located in Bolivia, 72 kilometers to the East from the capital of La Paz. Two fenced areas define the primary monuments, though recent estimates place the extension of the site up to 6km$^2$. The eight Tiwanaku site individuals originate from four different locations, and four different projects over thirty years. These individuals were on-site in storage facilities built by various efforts under the care of the local administration. Unfortunately, the information on their original context is not uniform. What is certain is that all these individuals originate from the within the ritual core: the base and revetments of the Akapana Platform (TW061, TW060), within the construction fill of the Pumapunku Platform (TW059), within the platform surrounding the Putuni courtyard (TW063), a collection of sacrifices and offerings in the area between Semi-Subterranean Temple and the Akapana Platform (TW097), and an assemblage of human remains and ceramics directly north of the Kalasasaya adjacent to a monolith (Monolito Descabezado) (TW056). Two individuals do not have a specific provenience (TW004, TW008) other than "Akapana" and "Museo Templo" respectively. The "Akapana" individual came from excavations on the Akapana during Bolivian led PAAK project in the mid-2000s, or the University of Chicago project of the 1990s. "Museo Templo" probably refers to the burials disturbed by the construction of the site museum directly outside the fenced monumental core on the South-West side of Akapana (Zone 1).

The individuals range to the period when Tiwanaku was a well-established center in the Titicaca Basin up to the period of abandonment. The well-excavated individuals (TW060, TW061) collected by the Bolivia project (PAAK) in 2004 and 2008, were located in the silt directly above the ground level of the base of the Akapana Platform. Producing dates centered around 950AD, these individuals are in a context with offerings similarly dated to the mid-10th century (Janusek, 2004). This context marks the period directly after the end of active construction and maintenance of the Akapana Platform (Vranich, 2001, Yaeger and Vranich, 2013).

Located slightly southeast between the Semi-Subterranean Temple and the Akapana Platform, individual TW097 (Proyecto Arqueológico Pumapunku-Akapana, 2006) is associated with an area of offerings set in silt above a prepared pebble plaza. These offerings align with the winter solstice sunset over the Kalasasaya Platform corner. TW056 (Monolito Descabezado) is adjacent to a stele whose iconographic style dates to the Late Formative Period (100 BC to 400 AD). The archaeologists interpret the four partial skeletons along with a substantial amount of decorated ceramics and other artefacts as the secondary deposit of ritual events. One skeleton was found facing down with upper and lower extremities flexed backwards, a position consistent with someone being bound.

The Putuni (TW063) individual found in the heavily looted southern exterior side of the platform. It would be difficult to place this skull in a more precise location other than the Putuni Platform. Individual from Pumapunku (TW059) was a full individual placed within the clean construction fill associated with the modification to the platform, explicitly raising the platform's height and the construction of a new set of hydraulics using reused ashlars.

**Lukurmata and Katari Valley**

Over a series of low hills to the North is the Katari valley, where the most significant site is Lukurmata (16°26’24”S 68°42’29”W). This site with a long occupation history has a monumental courtyard made with andesite ashlars repurposed from Tiwanaku. Extensive excavations by the University of Chicago project concentrated on the residential component and the terraforming for hydraulic and agricultural works. The individuals (TW013, TW020, TW027, TW028) are from burials associated with residences and span the period from before the rise of the Tiwanaku site to after its abandonment.

**Peru**

Samples from Peruvian individuals were obtained under permissions granted by Peruvian Ministerio de Cultura (formerly Instituto Nacional de Cultura).

**Maucallacta**

Maucallacta (15°41'05"S, 72°37'20"W) is located at the height of 3700 to 3800 masl.

It is a 62ha site with a central architectural complex of ca. 250 buildings, several platforms and plazas. The most prominent construction at the site is a large pyramidal structure with 30 x 24 m dimensions at base and 6 to 9.5 m tall and a huge platform with a 160 x 48 m plaza. Surrounding the main architectural sector of the site, there are five sectors comprising necropolis with numerous tombs. Maucallacta is considered the most important ceremonial and administrative center in the Coropuna and Solimana volcanoes area of the Inca Empire.

Individual CO066, analysed here, comes from a large collective tomb which was created by building up a space under a rock overhang. It is located in MA6 sector of the site at which five tombs have been found. Anthropological and paleopathological analysis of the bones form this site indicates that the inhabitants of Maucallacta were working quite hard physically. The equipment associated to the remains does not indicate elitism or foreign provenience of the people entombed there.

**Antaura**

The Antaura site occupies land located between around 3500 masl, on the eastern edge of the very narrow and deep valley of the Antaura River (The Maucallacta site is located on the opposite edge of the upper portion of the valley). It has been excavated under project “Condesuyos” ran by the Centre for Pre-Columbian Studies of the University of Warsaw and Universidad Catolica de Santa Maria in Arequipa since 1996. Antaura is a settlement complex made up of local architecture. These are small rectangular or oval buildings. The location of the buildings is rather chaotic, it extends along the edge of the valley. Inside the site there are some funeral constructions, located within the limits of the site on the north and north-east side. Despite a local character of the productive and housing complexes, in the sepulchral architecture there is a noticeable influence from Altiplano in form of oval-based chullpas rather than typical in this region rectangular ones.

Individual CO154 comes from a chullpa with multiple burials. DNA for the analysis was sampled from a femur.

**Culcunche**

The Culcunche-Quilluniyoc site is located at a part of a plateau limited by a small river called Río Blanco on the west side and by the edge of the plateau above Maucallacta. The site is located at a height of 4100 to 4150 masl and occupies an open plateau (*puna*), with pastoral (settlements) and funeral structures – space located on the culmination of the rocks that dominates the White River from the West. The central part forms the plaza, limited on two sides by the buildings (warehouses). The local complex combines warehouse and grazing area functions with regional routes.

The individual CO01was taken from the tomb of Beehive, located at the bottom of the rock cliff above the Rio Blanco River.

**Cotahuasi**

Individual CO193 was obtained from Museo Arqueológico “José María Morante Maldonado” de la UNSA in Arequipa. It is a mummy from a cliff overhang multiperson tomb found in the Cotahuasi Valley. The mummy comprised a woman and a young child wrapped together. CO193 DNA was extracted from the woman’s remains.

### 2. RADIOCARBON DATING

Radiocarbon dating was performed at the University of Waikato Radiocarbon Dating Laboratory and Poznan Radiocarbon Laboratory (Dataset S1*A*).

For age calibration we used SHCal13 curve (Hogg et al., 2013) in OxCal 4.3.2 (Bronk Ramsey, 2017) (Fig. S1). Additionally, due to air currents probably affecting the carbon concentrations by carrying the air from the Northern Hemisphere in the Titicaca region (Marsh et al., 2018) we calibrated the dates using mixed curve model (Dataset S1*A*). For all but one samples the median mixed curve ages were identical or up to 10 years apart. Three samples' median ages differed by up to 60 years between SHCal13 and mixed curve calibration. We used SHCal13 calibrated dates throughout the manuscript. The provided calibrated ages are rounded to the nearest 10.

**3. DNA EXTRACTION AND SEQUENCING**

All work with human remains, DNA extractions and library preparations were performed in facilities dedicated to work with ancient DNA. The laboratory was UV irradiated when not in use and all protocols recommended to prevent any contamination with modern DNA were followed. DNA was extracted from teeth or long bones. Samples were washed with ultrapure water and UV irradiated (245nm) for 10 min on each side in the laminar flow cabinet. Fragments of long bones and teeth were powdered in a cryogenic mill (SPEX CentriPrep). Additionally, in case of some teeth we tried drilling only cementum as it was described in Hansen et al., (2017). Extraction was performed following the protocol commonly used for retrieving short fragments of the ancient DNA (Dabney et al., 2013). Around 100-150 mg of powder was submerged in 1ml of extraction buffer (0.5 M EDTA pH = 8.0; 0.5% N-Laurylsarcosine; 0.2 mg Proteinase K) and digested overnight at 37°C. Supernatant was transferred to an assembly made up of MinElute silica spin column (Qiagen) mounted in the sterile Zymo-Spin reservoir and placed in the 50ml tube. Binding buffer (PB buffer, Qiagen) was added to supernatant in ratio 13:1. The solution was purified through the column spinning down at 1,500xg with slow acceleration for 6 minutes. Silica-membrane was washed twice with 750 µl of PE buffer (Qiagen). DNA was eluted with 60 µl of warmed EB buffer (Qiagen) (2x30ul). Extractions were performed for up to 15 samples accompanied with negative control.

Double-indexed sequencing libraries were constructed using 20 µl of DNA extract according to the protocol proposed by Meyer and Kircher (2010) with minor modifications: after blunt-end repair and fill-in enzymes were heat-inactivated for 20min at 75 °C and 80 °C, respectively. Every library possessed unique pair of P7 and P5 indexes. Number of cycles in indexing PCR was determined using qPCR with primers I7 and I8 (Meyer & Kircher, 2010). Indexing-PCR was performed using Pfu Turbo Cx DNA polymerase (Agilent) and 10 µl of DNA. Three independent PCRs were performed for each library to increase complexity. After amplifications they were pooled and SPRI purified. For a few samples we tried enriching endogenous DNA using myBaits WGE Human (Arbor Biosciences) according to the manufacturer protocol (Dataset S1*A*).

Sequencing libraries were pooled in equimolar ratios and sequenced on the Illumina platforms: either NextSeq550 platform (HighOutput, 75 cycles, single-end; MidOutput, 150cycles, paired-end), HiSeq4000 (50cycles, single-end) or NovaSeq600 (S1, 100cycles, single-end).

**4. DATA PROCESSING**

Sequencing reads were demultiplexed using bcl2fastq Conversion Software. Illumina adapters were trimmed and paired-end collapsed with AdapterRemoval v. 2 (Schubert et al., 2016). We used bwa mem to map sequencing reads to human reference genome h37db5 (Li & Durbin, 2010). Only reads with quality >30 and with length >30bp were retained. Duplications were removed using Samtools (Li et al., 2009).

Sequencing libraries were generated without USER-treatment. As it is known that deamination rate is the highest at the ends of the molecule, we trimmed 7nt from the beginning and from the end of the sequencing reads to minimize genotyping errors. For this purpose we used trimbam script from the bamUtils (Jun et al., 2015). In parallel, the reads without end-trimming were used to calculate the frequencies of the deamination at the 5' and 3' ends with mapDamage v.2 (Jónsson et al., 2013) to assess the authenticity of the obtained sequences as ancient DNA (Dataset S1*A).*

Mean genome coverage and mean fragment length were obtained with QualiMap (García-Alcalde et al., 2012) (Dataset S1*A*).

### 5. CONTAMINATION CONTROLS

Potential contamination was checked applying four different approaches. Three of them were based on calculation of heterozygosity rates on the haploid markers (mtDNA and X chromosome) as a measure of contamination. The first approach was schmutzi (Renaud et al., 2015) which estimates contamination in mtDNA using database of mitochondrial allele frequencies. We use parameters --notusepredC and –uselength. The second approach contamMix v1.0-10 (Fu et al., 2013) which estimates contaminant/authentic mixture proportions in mitochondrial sequence data for a set of potential contaminant genomes, in this case a panel of 311 worldwide mitogenomes as a potential contamination source was used. The second haploid marker available to check for contamination is the X-chromosome for male specimens. For that purpose we used ANGSD (Korneliussen et al., 2014) which in the first step calculates rate of heterozygosity on the each position and later estimates contamination using HapMap reference files with global allele frequency. We set minimum base quality to 30 and minimum mapping quality to 30 with other default parameters in the first step. Lastly, for the both females and males, contamination in the autosomes was measured using ContamLD version 1.0 which based on breakdown of the linkage disequilibrium (Nakatsuka et al., 2020a). For ContamLD analyses, default settings were used with PEL as reference population for all samples. All contamination results are reported in Dataset S1*A*. In case of three male samples (TW008, TW097 and TW098) the X-coverage was insufficient to estimate contamination (<200 SNPs available for analysis).

### 6. GENOTYPING

Genotyping was performed using the script pileupCaller - a part of sequenceTools (https://github.com/stschiff/sequenceTools), which randomly samples one read for every SNP position. Pseudohaplotypes of the analyzed individuals were called using list of 1240K SNPs (Haak et al., 2015) and we found that between 142,180 and 926,465 SNPs were overlapping with the 1240K set (Dataset S1A).

### 7. GENETIC AFFINITIES

Dataset from this study was merged with available genomic data for South America using program mergeit from the Admixtools package (Patterson et al., 2012). The data was merged with the available present-day and ancient genome data from South America (de la Fuente et al., 2018; Gnecchi-Ruscone et al., 2019; Lindo et al., 2018; Nakatsuka et al., 2020b; Posth et al., 2018; Reich et al., 2012). Because the present-day genotypes were obtained using various techniques, including shotgun sequencing and different Illumina microarrays, the final intersection gave 199,175 common SNPs. The number of SNPs intersected in the samples from this study ranged from 24,601 SNPs to 161,748 SNPs (Dataset S1_A). In downstream analyses we used published ancient genomes that possessed more than 20,000 SNPs intersected and were dated to the last 2000 years (Dataset S1*C*).

Principal Component Analysis (PCA) was computed using smartpca script from EIGENSOFT package (Patterson et al., 2006). We used 37 modern populations to compute PCs: Arara, Arhuaco, Ashaninka, Aymara, Aymara Ventilla, Bolivia Aymara, Cashibo, Chane, Chilote, Chono, Diaguita, Embera, Guahibo, Guarani, Huambisa, Hulliche, Inga, Jamamadi, Kaingang, Karitiana, Kogi, Parakana, Piapoco, Quechua, Shipibo, Surui, Ticuna, Titicaca Aymara, Titicaca Quechua, Titicaca Uros, Toba, Waunana, Wayuu, Wichi, Yaghan, Yanesha HighSelva, Yanesha IntermediateSelva (Dataset S1*C*). Using option lsqproject=YES and shrinkmode=YES options ancient individuals from this study and literature were projected.

Unsupervised admixture analysis was performed using software ADMIXTURE (Alexander et al., 2009). To convert data from the eigensoft format to PED format we used program convertf from the Admixtools package (Patterson et al., 2012). Prior to admixture analysis dataset was filtered and pruned with plink (Purcell et al., 2007). For admixture computation we used only individuals from this study and 288 published genomes from South America (Dataset S1*C*). This filtered dataset was firstly pruned for minor allele frequency below 0.01 (--maf 0.01) following with pruning for linkage disequilibrium between markers with a window size of 200, a step size of 5 and an $R^2$ threshold of 0.5 (--indep-pairwise 200 5 0.5). After all pruning activities 100,290 SNPs were kept. Every individual from this study met the software requirements (>10,000 SNPs) and was used in the analysis. Five replicates were done for each K (K=2 to K=15). The optimal K (K=5) was chosen based on the lowest cross-validation errors (Dataset S1*D*). Very similar admixture patterns in all Tiwanaku individuals were found for other K's with lower CV values (Fig. S2).

Outgroup $f_3$-statistics were estimated with qp3pop from ADMIXTOOLS (Patterson et al., 2012) in format *$f_3$(Ind, Test, Mbuti)*. Using completely divergent outgroup like Mbuti we were able to measure genetic drift between two individuals/population. In all calculations we ascribed each of the individuals analyzed in this study as '*Ind*' and every other individual/population from the dataset as '*Test*' (Dataset S1*C*). The higher $f_3$-values mean higher genetic affinity between two tested individuals (or individual-population). All obtained values were plotted using ggplot2 and forecats packages in R (Fig. S3). We created a dissimilarity matrix from $f_3$-statistics calculating the 1-$f_3$ values. Multidimensional scaling was performed using cmdscale from R package and the principal dimensions were plotted. Neighbor joining tree was generated using PHYLIP software (Felsenstein, 1993) and ancient Beringian individual *USA_USR1_AncientBeringian_1140BP.SG* was used as an outgroup (Moreno-Mayar et al., 2014).

The R package admixR (Petr et al., 2019) which utilizes the ADMIXTURE software suite was used to calculate $f_4$-statistics and for qpWave and qpAdm modeling. In all calculations option '*transversion only*' was applied. Standard errors were computed using jackknife block size: 0.050. We performed exhaustive calculations of $f_4$-statistics in the form *$f_4$(Mbuti, popTest; Ind1, Ind2*) and *$f_4$(Mbuti, Ind; groupTest1, groupTest2*) where *Ind* were individuals from this study while *popTest* or *groupTest* were a populations or the group of ancient individuals from the same region and period as defined in Nakatsuka et al., (2020b) or present-day populations (Dataset S1*C*). Significantly negative values of Z-score (Z > |3|) suggest that the studied *Ind1* shared more alleles with *Test* population than with *Ind2* (Dataset S2*A*) or with *Group1* than with *Group2* (Dataset S2*B*). To test if the individuals from Lukurmata are genetically similar we calculated $f_4$-statistics in the form *$f_4$(Mbuti, Test; LUKind1, LUKind2)* where *LUKind1* and *LUKind2* were all possible pairs of Lukurmata individuals iterating over groups (*Test*) formed of ancient individuals from the same period and region as defined in Nakatsuka et al. (2020b) (Dataset S2*C*). Further, genetic homogeneity of Lukurmata individuals with *ancientTiticaca* group (*RioUncallane, Bolivia_Miraflores_1100BP_MH, Bolivia_Iroco_1050BP_MH*) were tested using $f_4$-statistics in the form *$f_4$(Mbuti, Test; LUK, ancientTiticaca)* (Dataset S2*D*).

To investigate genetic affinity between each individual from the ritual core of Tiwanaku (TW) with respect to the *ancientTiticaca* group we performed statistics of the form *$f_4$(Mbuti, ancTiticaca; TW1, TW2).* We tested whether each individual from Lukurmata and Tiwanaku ritual core shares more alleles with either ancient Peru group (*NorthernPeruHighlands, NorthernPeruCoast, CentralPeruCoast, SouthernPeruHighlands, SouthernPeruCoast* as defined in Nakatsuka et al., (2020b)) or with *ancientTiticaca* by computing $f_4$-statistics in the form *$f_4$(Mbuti, ancPeru; Ind, ancTiticaca)* (Dataset S2*F*). The significant affinity of the TW056 individual to Amazonian Cashibo group was tested with *$f_4$ (Mbuti, Cash; TIW, ancTiticaca)* (Dataset S2*G*).

We used qpWave (version: 600) to determine the minimum number of ancestry sources for either of the individuals or group from this study (Lukurmata and Tiwanaku). As a left (‘*Source*’) population we used groups of ancient individuals defined by Nakatsuka et al. (2020b) – all ancient Peruvian, ancient NorthChile and ancient Titicaca as well as and present-day populations from Peruvian and Brazilian Amazonia, Colombia and GranChaco (Dataset S1*C*). To set right (‘*Reference*’) set we applied ‘rotating’ strategy suggested by Harney et al., (2020). It means that to the basic set of reference populations that include *Argentina_ArroyoSeco2_7700BP*, *Peru_Cuncaicha_4200BP*, *Peru_Lauricocha_3500BP*, *Brazil_LapaDoSanto_9600BP*, *Wichi* we added also populations from the ‘source’ list that were not used as a ‘source’. Under this ‘rotating’ approach populations are consistently moved from the set of ‘*Source*’ to set of ‘*Reference*’ populations.

The same strategy has been applied for qpAdm modeling (qpAdm version: 1000, allsnps: YES). We started modeling from the single source of ancestry followed by models with two- and three-sources of ancestry (Dataset S3).

### 8. MITOCHONDRIAL DNA AND Y-CHROMOSOME HAPLOTYPING

For every studied individual we were able to obtain complete sequence of mitochondrial genome. Sequencing reads were mapped to reference mtDNA sequence (rCRS, NC_012920) using bwa mem and several programs from Samtools (Li et al., 2009) were used to remove sequence shorter than 30 bp, with mapping quality under 30 and representing duplicates. Variations and consensus sequences were called using bcftools (Li et al., 2009). Bam files were checked manually using Tablet (Milne et al., 2013) and only positions with 3x coverage were accepted. Nucleotide positions 309-311, 523-524, 3107, 8271-8279, 16182, 16183 were masked and omitted in haplogroup calls. Using Haplogrep2 (Phylotree 17) (Weissensteiner et al., 2016) every mtDNA sequences was assigned to one of the main South American haplogroups (Dataset S1*A*). Y-chromosome haplogroups were determined using Yleaf software v2.1 (Ralf et al., 2018) (Dataset S1*A*).

**Fig. S1**

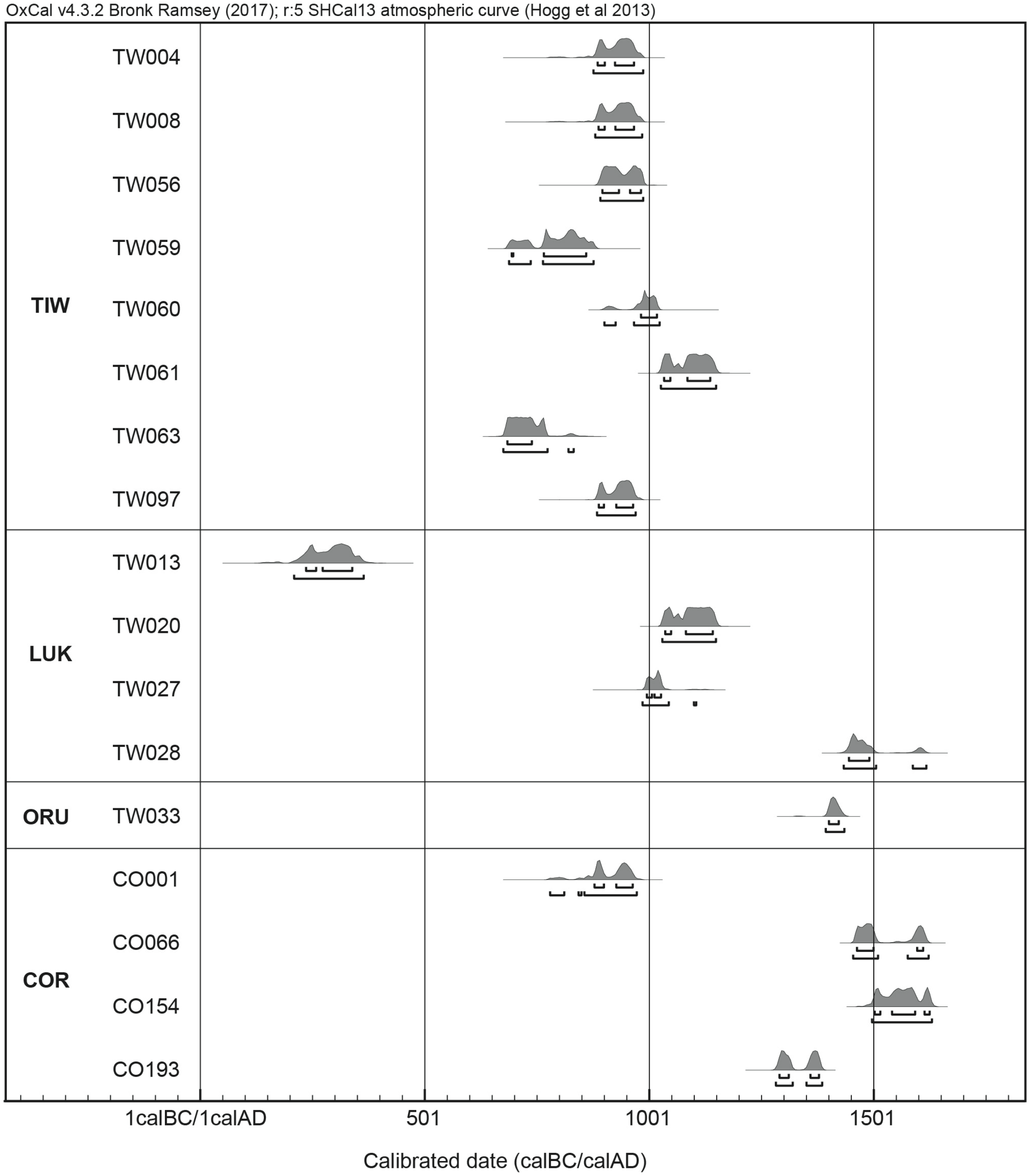


**Fig. S1** Probability distribution plots for the SHCal13 (Hogg et al. 2013) calibrated dates for all individuals analyzed in the study. Calibration was obtained using OxCal v4.3.2 (Bronk Ramsey 2017). Brackets underneath the plots delimit 68.2% and 95.4% probability, respectively.

### Fig. S2

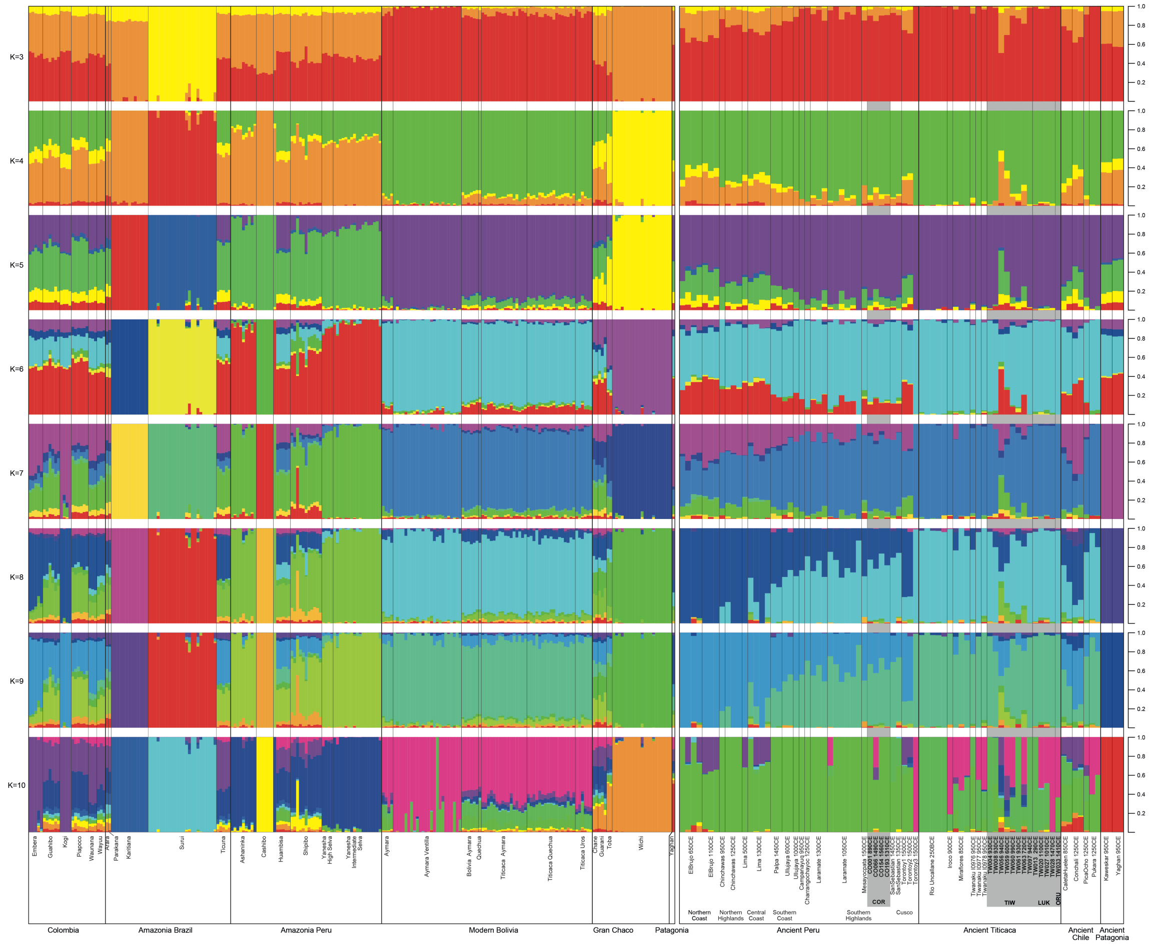


**Fig. S2.** ADMIXTURE plots for K=3 to K=10. Individuals order identical to that in Figure 2C; modern individuals in left panel and ancient individuals in right panel.

**Fig. S3.**

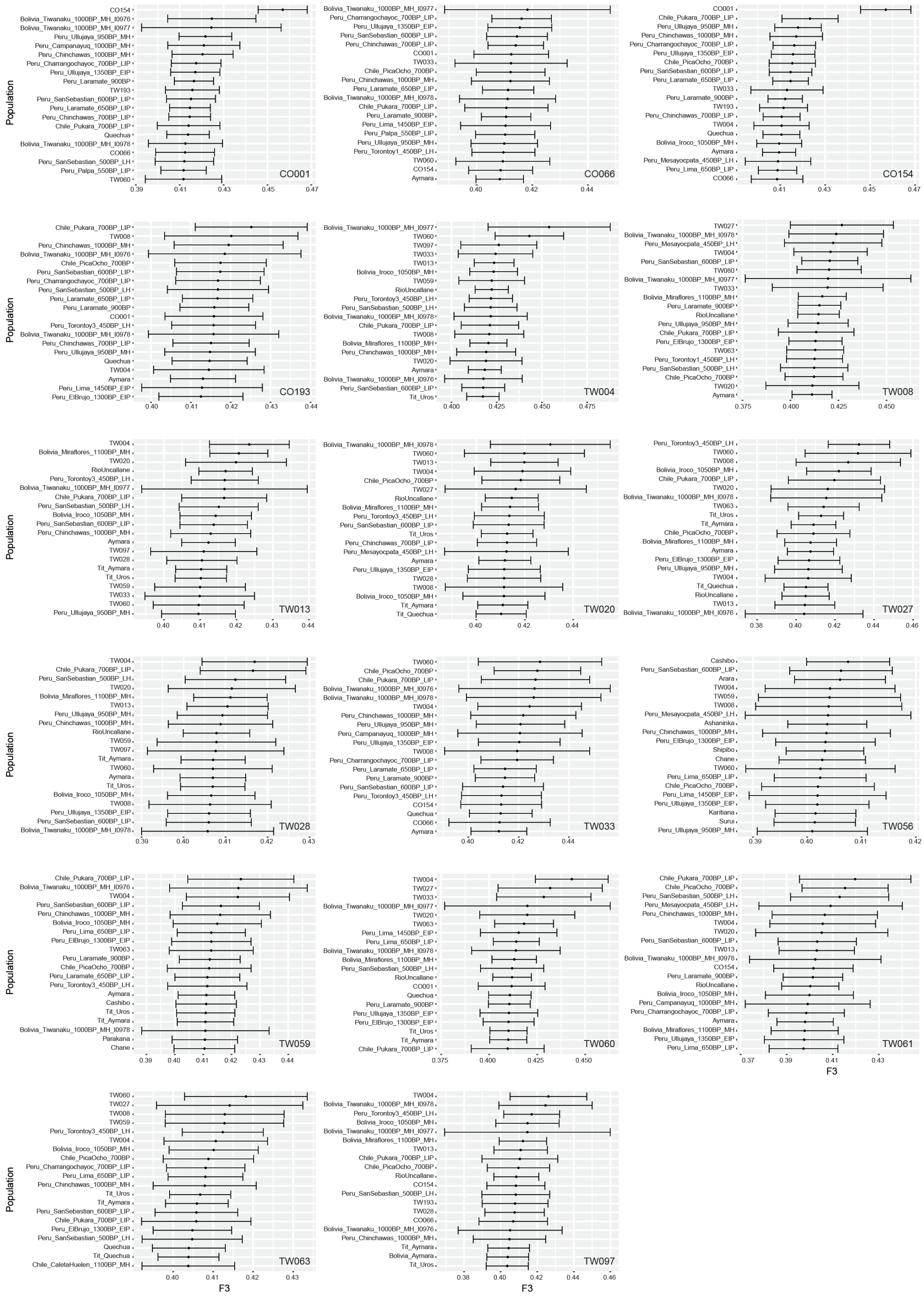


**Fig. S3.** Outgroup $f_3$-statistics for each individual from this study.

Dataset S1 (separate file). Sample and sequencing information, dataset of published genomes used for downstream analyses, admixture CV values.

Dataset S2 (separate file). $f_4$-statistics.

Dataset S3 (separate file). qpWave and qpAdm analyses.
